# Case-control design identifies ecological drivers of endemic coral diseases

**DOI:** 10.1101/662320

**Authors:** Jamie M. Caldwell, Greta Aeby, Scott F. Heron, Megan J. Donahue

**Affiliations:** Hawaii Institute of Marine Biology, University of Hawaii at Manoa, Hawaii, USA; ARC Centre of Excellence for Coral Reef Studies, James Cook University, Townsville, Australia; Qatar University, Department of Biological & Environmental Sciences, Doha, Qatar; Marine Geophysical Laboratory, Physics, College of Science and Engineering, James Cook University, Townsville, Australia; NOAA Coral Reef Watch, College Park, Maryland, USA

## Abstract

Endemic disease transmission is an important ecological process that is challenging to study because of low occurrence rates. Here, we investigate the ecological drivers of two coral diseases -- growth anomalies and tissue loss -- affecting five coral species. We first show that a statistical framework called the case-control study design, commonly used in epidemiology but rarely applied to ecology, provided high predictive accuracy (67-82%) and disease detection rates (60-83%) compared with a traditional statistical approach that yielded high accuracy (98-100%) but low disease detection rates (0-17%). Using this framework, we found evidence that 1) larger corals have higher disease risk; 2) shallow reefs with low herbivorous fish abundance, limited water motion, and located adjacent to watersheds with high fertilizer and pesticide runoff promote low levels of growth anomalies, a chronic coral disease; and 3) wave exposure, stream exposure, depth, and low thermal stress are associated with tissue loss disease risk during interepidemic periods. Variation in risk factors across host-disease pairs suggests that either different pathogens cause the same gross lesions in different species or that the same disease may arise in different species under different ecological conditions.

## Introduction

Disease is an ecologically and evolutionarily important process in shaping populations, communities, and ecosystems [1–5], but identifying factors that promote disease at low endemic levels or between epidemics is challenging because disease occurrences are rare in space and time. Disease can affect populations, communities, and ecosystems through multiple pathways, such as changing the physiology, behavior, distribution, abundance, and fitness of organisms (e.g., [3,6,7]). For example, when the trematode *Plagioporous sp.* infects the coral *Porites compressa* it reduces coral growth and causes infected polyps to appear as bright swollen nodules that cannot retract into their skeletal cups, thus increasing fish predation and affecting coral physiology, behavior, and fitness [8]. As another example, infection in three-spined sticklebacks by the fish parasite *Gyrodactylus* spp. has been shown to alter the zooplankton community structure and nutrient cycling in streams and lakes, which in turn, affect the survival and fitness of the subsequent fish generation [5]. Despite the ecological and evolutionary importance of disease, understanding the factors that increase individual disease risk for endemic diseases with low prevalence or during interepidemic periods for diseases with epidemic cycles can be challenging because of the low probability of observing diseased individuals during non-outbreak periods.

With limited occurrence data relative to nonoccurrence data, commonly used biostatistics such as logistic regression can significantly underestimate the probability of rare events such as disease occurrence [9]. Further, such models are likely to have high predictive accuracy (proportion of correctly identified event and nonevent observations), which can be misleading: for example, if 3 of 100 individuals are diseased, a model where all observations are predicted healthy would return 97% predictive accuracy but provide very limited information about disease occurrence or causation. Ecologists interested in rare events such as insect outbreaks and species invasions could benefit from adopting statistical approaches from fields of inquiry concerned with rare phenomena. One statistical framework that works well with low event-to-nonevent ratio datasets is called the case-control study design [10]. The case-control study design is commonly used in epidemiology where the number of people affected by a condition is very small relative to the potential control population (e.g., few people are HIV+ compared with the general population). In the case-control design, for each subject with some condition, a control subject is selected that does not have the condition but is otherwise similar in terms of factors such as age, sex, and occupation. This approach naturally lends itself to many ecological studies with low event-to-nonevent ratios such as wildlife disease occurrence, with some modifications. In particular, the case-control design is typically applied to cohorts of individuals who are followed through time. For wildlife diseases, cohort data are rarely available. Instead, we propose that wildlife studies can apply the case-control design by retrospectively sub-sampling cross-sectional data that are widely available.

Here, we compare logistic regression models with and without a case-control design (based on retrospective sub-sampling) to answer three ecological questions for marine wildlife diseases: 1) how accurately can we predict disease occurrence; 2) what are the ecological factors associated with maintaining low levels of disease occurrence; and 3) are risk factors more similar for closely related species? We used coral diseases for this case study because (i) they can be extremely rare (<0.1% of disease occurrence observations); (ii) lesions can be visually identified; (iii) they can cause high mortality or strongly reduce fecundity with likely long-term population effects; and (iv) large datasets are publicly available on coral health observations and hypothesized risk factors.

In this study, we investigate two types of coral diseases – growth anomalies and tissue loss diseases – affecting five species of coral from two widely distributed genera in the Indo-Pacific Ocean. Growth anomalies are chronic, protuberant masses of coral skeletons (i.e., tumors) that reduce growth, fecundity, and survival [11–14]. Growth anomalies in some coral genera (e.g. *Porites* spp.) have been associated with larger coral colonies [15] located in shallow water (<3 m) [12] with elevated nutrients [16] and in regions with high human populations [17]. Tissue loss diseases, or white syndromes, usually have very low endemic prevalence levels (<1% in the Indo-Pacific) [18] but can be associated with rapid outbreak events resulting in localized mass mortality [19–23]. This suite of diseases is characterized by progressive tissue loss across the coral colony with lesions progressing slowly (chronic to subacute) or rapidly (acute). Bacterial pathogens cause many tissue loss diseases [24–27] and can be associated with high coral cover (i.e., density dependent transmission) and temperature stress (e.g. anomalously warm or cold ocean temperatures) [28–32]. Based on these previous findings, we hypothesize that long-term, chronic stressors are more likely to drive growth anomaly persistence whereas locations with pulses of short-term, acute stressors are more likely to be associated with interepidemic tissue loss occurrence. Growth anomalies and tissue loss appear on numerous coral species and likely constitute multiple diseases. Thus, we investigate each host-disease pair separately.

## Results

### Comparison between statistical approaches

Logistic regression models with and without a case-control design had high overall predictive accuracy (proportion of correctly predicted healthy and disease observations) for all host-disease pairs, but only the models with the case-control design accurately predicted disease occurrence (proportion of disease observations the model correctly predicted as diseased; i.e., the true positive rate). Using colony health state as a binary response variable (0 indicating healthy, 1 indicating diseased; Table 1) and a suite of hypothesized risk factors (Tables 2-3), we found that mean predictive accuracies ranged from 98-100% for logistic regression without the case-control design and 67-82% for logistic regression with the case-control design (Fig. 1). True positive rate, or the proportion of disease observations the model correctly predicted as diseased, is of greatest interest for rare event prediction and mean values ranged from 0-17% for logistic regression without the case-control design and 60-83% for logistic regression with the case-control design (Fig. 1). Given the moderate-to-high mean predictive accuracies (67-82%) and mean true positive rates (60-83%) for the case-control design, we report only results using the case-control design below.

**Figure 1:**
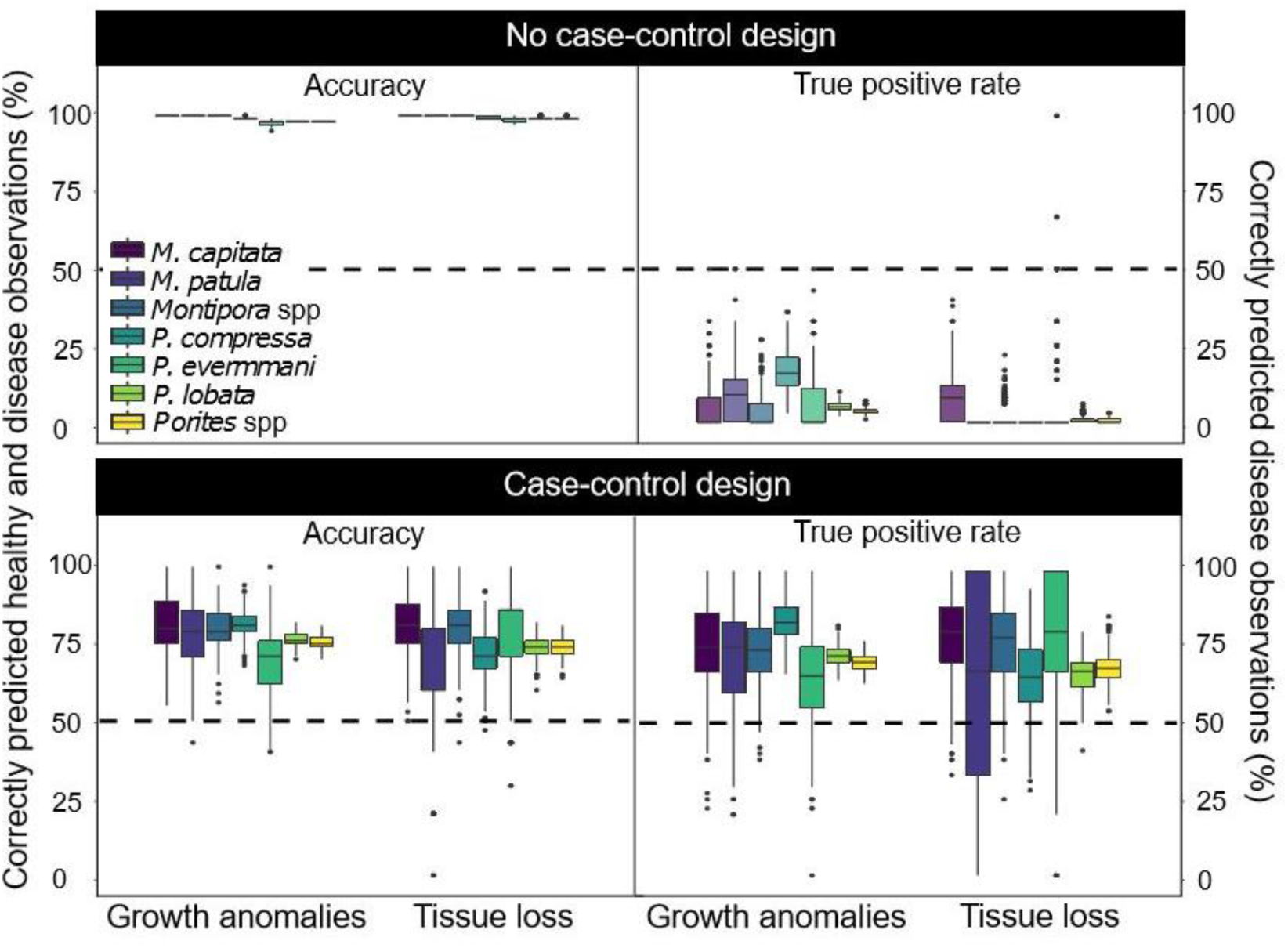
Logistic regression with case-control design has higher disease detection rates compared with logistic regression without case-control design. Boxplots of accuracy (percent correctly predicted healthy and disease observations) and true positive rates (percent correctly predicted disease observations) across 500 randomly sampled test datasets (see Methods) for each host-disease pair included in this study. The top panel shows results for logistic regression models without the case-control design and the bottom panel shows results for logistic regression models with the case-control design. The left side of the panels show accuracy results and the right side of the panels show true positive rates. Boxplots are grouped by disease type and colored by host; models at the genus level include observations from all hosts species within that genus (e.g., *Montipora* spp. includes all observations from *Montipora capitata* and *Montipora patula*).

**Table 1:** Healthy observations far exceeded disease observations for all host-disease pairs. Each row corresponds to the total number of disease and healthy colony observations used in this analysis for a specific disease (GA=growth anomalies, TL=tissue loss) and host (species or genus) pair. Hosts at the genus level include observations from all hosts species within that genus (e.g., *Montipora* spp. includes all observations from *Montipora capitata* and *Montipora patula*). In the analysis, we used 80% of observations (training data) to create the models and the remaining 20% of observations (test data) for model validation. We calculated accuracy and true positive rates based on the test data. There was an equal number of disease observations in the test data for logistic regression with and without the case-control design. In contrast, the number of healthy observations differed between the test data for logistic regression with and without the case control design: in the case-control design, the number of healthy observations was equal to the number of disease observations whereas without the case-control design the number of healthy observations in the test data was equal to ∼20% of the total number of healthy observations.

| Disease type | Host | Total disease | Total healthy |
| --- | --- | --- | --- |
| GA | *Montipora capitata* | 49 | 35864 |
| GA | *Montipora patula* | 35 | 23118 |
| GA | *Montipora* spp. | 84 | 58982 |
| GA | *Porites compressa* | 160 | 17889 |
| GA | *Porites evermanni* | 42 | 1779 |
| GA | *Porites lobata* | 1396 | 64312 |
| GA | *Porites* spp. | 1598 | 83980 |
| TL | *Montipora capitata* | 39 | 28669 |
| TL | *Montipora patula* | 12 | 17480 |
| TL | *Montipora* spp. | 51 | 46149 |
| TL | *Porites compressa* | 89 | 16915 |
| TL | *Porites evermanni* | 17 | 1567 |
| TL | *Porites lobata* | 351 | 61489 |
| TL | *Porites* spp. | 457 | 79971 |

**Table 2.** Risk factors (predictor variables) included in candidate models for each disease type with the direction and mechanism(s) of hypothesized relationships. Relationships and their proposed mechanisms described previously in the literature are referenced whereas hypothesized relationships based on personal observations are not referenced. References followed by an asterisk indicate relationships found for other type(s) of coral disease and/or related risk factors.

| Risk factor | Hypothesized relationship with disease risk | Hypothesized mechanism(s) relating risk factor to disease risk | Disease Type |
| --- | --- | --- | --- |
| Colony size | Positive | • Larger corals have more tissue exposed to pathogens<br>• Larger, older corals have a longer duration of exposure to pathogens/stress<br>• Larger, older corals have reduced immune capacity | Growth anomalies [15]<br>Tissue loss [33] |
| Host density | Positive<br>Negative | • Disease spread increases among many, closely spaced colonies<br>• Disease spread is reduced in high density reefs because colonies are smaller | Growth anomalies [17]<br>Tissue loss [33,56] |
| Depth | Negative | • Multiple environmental conditions become less stressful to coral with increasing depth: deeper water absorbs more light, has cooler temperatures, and fewer re-suspended sediments | Growth anomalies [12,29]<br>Tissue loss [29,56] |
| Water temperature (Hot Snap) | Positive<br>Negative | • High temperature reduces coral immune capacity<br>• High temperature increases pathogen survival, growth rate, and/or virulence<br>• High temperature reduces pathogen survival, growth rate, and/or virulence | Growth anomalies [29,57]<br>Tissue loss [28–30] |
| Human population size | Positive | • People increase reef degradation and physical damage to coral | Growth anomalies [17,58]<br>Tissue loss [59] |
| Embayment | Positive | • Areas inside bays have poor water quality (e.g., higher levels of nitrogen, sediments, and land-based pollutants) and are more likely to retain pathogens | Growth anomalies [15]<br>Tissue loss [19] |
| Herbivore fish abundance | Positive<br>Negative | • Fish alter algae composition to favor algal species that harbor pathogens<br>• Fish spread pathogens via defecation on corals<br>• Fish reduce macroalgae that harbor pathogens<br>• Fewer fish indicate higher fishing intensity and associated impacts | Growth anomalies Tissue loss [18]<br>[60–63]* |
| Agriculture and | Positive | • Pesticides and fertilizers increase coral stress and susceptibility to infection | Growth anomalies |
| golf course runoff |  |  | [41,64]* |
| Irradiance | Positive | • Light increases oxidative stress in coral | Growth anomalies [58] |
| Wave energy (chronic) | Negative | • Locations with high water flow reduce temperature and light stress to coral | Growth anomalies [57] |
| Wave energy (acute) | Positive<br>Negative | • Pulses of water flow increase contact rate between corals and pathogens<br>• Pulses of water flow limit duration of coral exposure to pathogens<br>• Pulses of water flow increase coral abrasion via sediment or physical damage | Tissue loss |
| Stream exposure | Positive | • Stream runoff increases contact rate between corals and pathogens<br>• Stream runoff reduces coral immune capacity by decreasing salinity and increasing sedimentation and exposure to land-based pollutants | Tissue loss [65] |
| Rainfall anomaly | Positive | • Rainfall reduces coral immune capacity by decreasing salinity and increasing land-based runoff of pathogens or pollutants<br>• Rainfall increases contact rate between corals and pathogens | Tissue loss [66]* |
| Chlorophyll-a (proxy for nutrients and phytoplankton) | Positive | • High chlorophyll-a concentration increases coral stress and susceptibility to infection<br>• High chlorophyll-a concentration leads to increased zooplankton, which can serve as disease vectors | Tissue loss [18] |

**Table 3:** Characterization of biological, environmental, physical, and anthropogenic risk factors (predictor variables) used in candidate models.

| Risk factor | Ecological category | Characteristic | Description | Minimum value | Maximum value | Source |
| --- | --- | --- | --- | --- | --- | --- |
| Colony size | Biological | Site | Maximum diameter or average size of size class (cm). | <1 | 300 | Survey data<sup>A</sup> |
| Host density | Biological | Site | Number of colonies divided by survey area (m<sup>-2</sup>). | 0.02 | 9.49 | Survey data<sup>A</sup> |
| Herbivore fish abundance | Biological | Site | Pooled estimate of herbivorous fish calculated between 2012 and 2013 (g m<sup>-2</sup>). | 0.06 | 0.42 | [67] |
| Chlorophyll-a | Environmental | Chronic | Maximum anomaly calculated within a 30-meter boundary of the coast between 2003 and 2013 from NASA MODIS satellite imagery (mg m<sup>-3</sup>). | 0.0004 | 0.3566 | [68]<sup>B</sup> |
| Irradiance | Environmental | Chronic | Photosynthetically active radiation (PAR) average annual frequency of anomalies calculated within a 30-meter boundary of the coast between 2003 and 2013 from NASA MODIS satellite imagery (mol m<sup>-2</sup> day<sup>-1</sup>). | 0.13 | 0.27 | [68]<sup>B</sup> |
| Hot Snap | Environmental | Acute | Magnitude and duration of heat stress above a summertime threshold at 5 km resolution (°C-week). | 0 | 3.6 | NOAA CRW<sup>C</sup> |
| Rainfall | Environmental | Acute | Monthly rainfall anomaly interpolated from rain gauge data, expert knowledge, radar observations, meteorological model simulations and vegetation data. Anomalies were calculated as the rainfall variation in any given month and year from the 1978-2007 mean. | 0.04 | 4.35 | [69]<sup>D</sup> |
| Depth | Physical | Site | Distance below sea surface (m). | 0.6 | 26.0 | Survey data<sup>A</sup> |
| Embayment | Physical | Site | Binary variable indicating whether survey site is outside or inside of semi-enclosed habitat. | 0 | 1 | Calculated in this study |
| Stream exposure | Physical | Acute | Inverse planar distance between survey site and stream mouth in nearest watershed (m<sup>-1</sup>). | 2.72 | 3.33 | Calculated in this study |
| Wave energy | Physical | Acute | Average annual maximum anomaly of wave energy flux calculated between 2000 and 2013 (kW m<sup>-1</sup>). | 0.08 | 285.74 | [68]<sup>B</sup> |
| Wave energy | Physical | Chronic | Average annual maximum monthly mean of wave energy flux calculated between 2000 and 2013 (kW m<sup>-1</sup>). | 0.04 | 103.48 | [68]<sup>B</sup> |
| Agricultural and golf | Anthropogenic | Chronic | Proxy for fertilizer and chemicals like pesticides and herbicides discharge from agricultural land and golf | 0.00 | 0.53 | [70] |
| course runoff |  |  | courses calculated as area of cultivated land and golf course per watershed (km<sup>2</sup>) with a distance-decay function. |  |  |  |
| Human population size | Anthropogenic | Chornic | Number of people living within 1 km of coastline nearest to survey site. | 0 | 4425 | NASA SEDAC GPW v4<sup>E</sup> |
| Public datasets:<br>A. Hawai'i Coral Disease database: https://www.sciencedirect.com/science/article/pii/S2352340916304607<br>B. Ocean Tipping Points: http://www.pacioos.hawaii.edu/projects/oceantippingpoints/#data<br>C. NOAA Coral Reef Watch CoralTemp: https://coralreefwatch.noaa.gov/satellite/coraltemp.php<br>D. Rainfall Atlas of Hawai'i: http://rainfall.geography.hawaii.edu/<br>E. NASA SEDAC Gridded Population of the world v4: http://sedac.ciesin.columbia.edu/data/collection/gpw-v4 |  |  |  |  |  |  |

### Ecological risk factors of different host-disease pairs

Of the 19 risk factors (predictor variables) we initially examined (see Methods), 14 were included in our candidate models (with the risk factors differing by disease type; Table 2), but the most predictive risk factors varied by host-disease pair. The 14 risk factors can be broadly classified into four categories: biological (colony size, host density, and herbivorous fish abundance), environmental (chlorophyll-*a*, irradiance, Hot Snap, and rainfall), physical (depth, embayment, acute and chronic wave exposure, and stream exposure), and anthropogenic (agricultural and golf course runoff and human population size).

All the risk factors that we hypothesized could be associated with growth anomalies were identified as important for at least one host-disease pair, with more similar risk factors associated with more closely related hosts. We found one biological risk factor (colony size; Fig. 2) common to all hosts, with an approximate 36-137% increase in disease risk of for every 1 cm increase in colony size above the host specific average size, given all other risk factors are held at their average values (Table 4). Herbivore fish abundance was the next most common risk factor, with an approximate 5-15% decrease in disease risk for every gram increase in herbivorous fish per meter squared (Table 4). *Montipora* species were positively associated with size, and negatively associated with herbivore fish abundance (*M.* capitata) or host density (*M. patula*); the genus model (*Montipora* spp.) consisted of all three biological risk factors (Table 4). In contrast, the *Porites* genus model (*Porites* spp.) only partially overlapped with the risk factors identified in the species-specific models (Table 4). *P. compressa* was associated with all risk factors other than irradiance, whereas *P. evermanni* was only associated with irradiance (in addition to colony size) (Table 4). *P. lobata* was associated with biological (host density and herbivore fish abundance), environmental (Hot Snap), physical (depth), and anthropogenic (agriculture and golf course runoff) risk factors, and most closely overlapped with the genus model (Table 4), likely because the majority of *Porites* observations were observations of *P. lobata* (Table 1). Agriculture and golf course runoff increased disease risk for *P. lobata* and *P. compressa* by approximately 23 and 80%, respectively, for every km^2^ increase in cultivated land.

**Figure 2:**
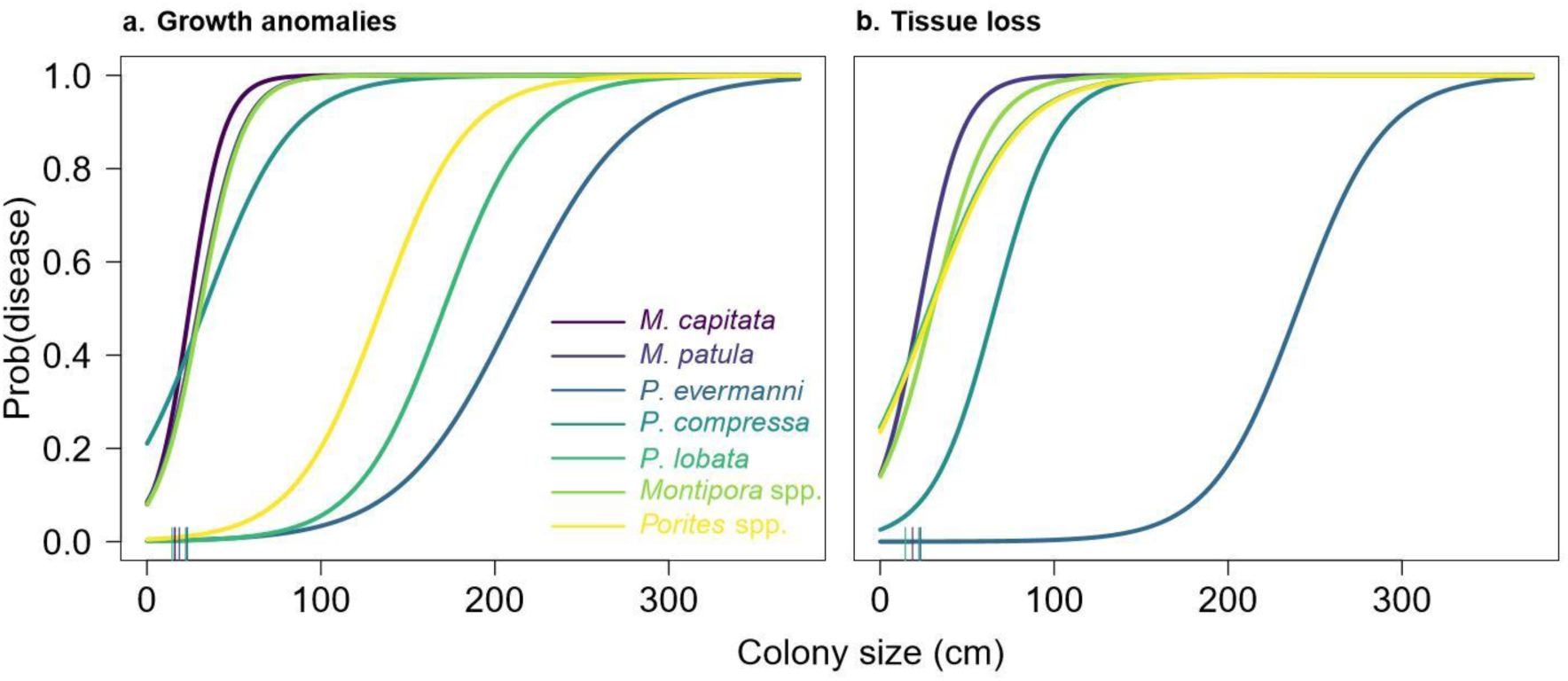
Larger colonies increase disease risk for most host-disease pairs. Lines indicate the expected probability of disease occurrence given colony size for a) growth anomalies and b) tissue loss. *M. capitata* is not shown in b because colony size is not a risk factor in the *M. capitata* tissue loss model (Table 5). Lines are colored by host species. Using the same color scheme, average colony size is indicated along the horizontal axis (*Montipora capitata* = 16 cm; *Montipora patula* = 19 cm; *Porites compressa* = 23 cm; *Porites evermanni* = 22 cm; *Porites lobata* = 15 cm). Fig. S1 shows individual plots for each host-disease pair (excluding genus models for visibility) with data points overlaid for one randomly sampled case-control training dataset.

**Table 4:** All hypothesized risk factors were associated with at least one host-disease pair for growth anomalies. Average coefficient values based on model runs across 500 case-control datasets for all host-disease pairs. Coefficient values indicate the magnitude and direction for the expected change in disease risk given a one-unit change in the risk factor, holding all other risk factors at their average values. Since the slope of the logistic curve is steepest at its center, coefficient values can be divided by 4 (“divide by 4 rule”) to reasonably approximate the upper bound of the predicted difference in disease risk corresponding to a one-unit difference in the risk factor [71].

| Host | Intercept | Size | Density | Fish | Hot Snap | PAR | Depth | Bay | Wave exposure | Ag & Golf | Human pop. |
| --- | --- | --- | --- | --- | --- | --- | --- | --- | --- | --- | --- |
| *M. capitata* | 1.92 | 5.46 |  | -0.60 |  |  |  |  |  |  |  |
| *M. patula* | 0.49 | 2.40 | -0.59 |  |  |  |  |  |  |  |  |
| *Montipora spp.* | 1.00 | 3.65 | -0.50 | -0.47 |  |  |  |  |  |  |  |
| *P. compressa* | -0.82 | 1.22 | -1.64 | -0.26 | -5.54 |  | 0.65 | 3.07 | -0.80 | 3.18 | -0.42 |
| *P. evermanni* | 4.05 | 2.39 |  |  |  | -1.93 |  |  |  |  |  |
| *P. lobata* | -0.58 | 1.34 |  | -0.19 | -6.99 |  | -0.62 |  |  | 0.93 |  |
| *Porites spp.* | 0.56 | 1.43 | 0.13 | -0.24 | -6.31 |  | -0.50 |  |  |  |  |

**Table 5:** Tissue loss associated with different biological, environmental, physical, and anthropogenic risk factors across host-disease pairs. Average coefficient values based on model runs across 500 case-control datasets for all host-disease pairs. Coefficient values indicate the magnitude and direction for the expected change in disease risk given a one-unit change in the risk factor, holding all other risk factors at their average values. Since the slope of the logistic curve is steepest at its center, coefficient values can be divided by 4 (“divide by 4 rule”) to reasonably approximate the upper bound of the predicted difference in disease risk corresponding to a unit difference in the risk factor [71].

| Host | Intercept | Size | Density | Fish | Hot Snap | Depth | Stream exposure | Wave exposure | Human pop. |
| --- | --- | --- | --- | --- | --- | --- | --- | --- | --- |
| *M. capitata* | -0.18 |  |  | -1.96 |  | -1.48 | -1.77 | -1.40 |  |
| *M. patula* | 0.69 | 2.53 |  |  |  |  |  |  |  |
| *Montipora spp.* | 0.60 | 2.71 |  | -1.28 |  |  | -0.77 | -0.76 |  |
| *P. compressa* | -2.37 | 1.45 |  |  | -3.49 |  | 0.43 |  |  |
| *P. evermanni* | 6.84 | 4.07 |  |  |  | -1.52 |  | -1.34 | -2.01 |
| *P. lobata* | -8.45 | 1.32 | -0.39 |  | -1.42 | -0.23 |  | 0.22 | -0.25 |
| *Porites spp.* | -3.69 | 1.42 | -0.29 |  | -4.71 | -0.23 |  | 0.16 | -0.27 |

For tissue loss diseases, we identified eight hypothesized risk factors associated with disease occurrence (five associated with *Montipora* species and seven associated with *Porites* species) (Table 5). Like growth anomalies, we found a positive association with colony size for most hosts with an approximate 33-102% increase in disease risk for every 1 cm increase in colony size (Table 5, Fig. 2). We found no overlap in risk factors for *Montipora capitata* and *Montipora patula*: *M. capitata* was associated with one biological (herbivore fish abundance) and three physical (depth, stream exposure, and wave exposure) risk factors whereas *M. patula* was only associated with one biological (colony size) risk factor (Table 5). Further, all risk factors for *M. capitata* had strong negative effects (Table 5). We identified the exact same risk factors for the *Porites* genus model and the *P. lobata* model (Table 5), which included biological (colony size, host density), environmental (Hot Snap), physical (depth, wave exposure), and anthropogenic (human population size) risk factors. However, Hot Snap had a stronger effect in the genus model compared with the *P. lobata* model, potentially indicating differences in thermal sensitivities across morphologies, as the coefficients for Hot Snap in the genus model and the *P. compressa* model were similar (Table 5). *P. evermanni* overlapped with four of those risk factors (colony size, depth, wave exposure, and human population size). However, the effect of wave exposure differed by host: higher wave exposure was associated with an increased risk of tissue loss in *Porites* spp. and *P. lobata* but reduced risk of tissue loss in *P. evermanni* (Table 5). The *P. compressa* model was the most distinct of the *Porites* models, with associations with colony size, Hot Snap, and stream exposure (Table 5).

## Discussion

Understanding the ecological conditions that promote endemic diseases with low prevalence and for epidemic diseases during interepidemic periods is central for understanding disease dynamics but can be difficult to investigate using common biostatistical approaches. In this study, we adopted statistical methods better suited towards highly skewed occurrence data to investigate ecological correlates of several coral diseases. We show that the case-control study design improved our ability to predict disease and healthy colony observations with relatively high specificity given a suite of ecological conditions. In contrast, a more traditional biostatistical approach yielded high overall predictive accuracy (heavily biased by successful predictions of healthy state) but often failed to accurately predict the events of interest: disease occurrences. This study highlights the usefulness of the case-control design for ecological questions using infrequent occurrence data. In this study, we identified biological, environmental, physical, and anthropogenic drivers that promote low levels of coral disease and explored how those factors differ across host-disease pairs. These results provide insight into factors that should be explored further through experimental and observational studies and that may be favorable for intervention strategies.

Among the suite of ecological risk factors that we included in our study, we found that disease risk was consistently higher for larger colonies across host-disease pairs. A positive size-disease relationship has been found in other coral disease studies [15,33,34] and could be associated with exposure area or duration of exposure to potential disease causing agents (biotic or abiotic), or immune function, which is hypothesized to decline with age [35]. Further, a positive relationship between disease prevalence and coral cover has been well established for infectious diseases like tissue loss [13,28,30,31]. However, because the relationship between coral size-frequency distributions and coral cover is not straightforward [36, 37], it is unknown whether disease prevalence is higher in locations with high coral cover because of many small colonies (many potential susceptible hosts and/or reservoir species) or fewer large colonies (higher susceptibility of certain individuals). Our results provide evidence that highly susceptible corals (i.e., large colonies) drive the positive relationship between disease prevalence and coral cover. Although we found a positive linear relationship between disease risk and size within the range of 0 to 300 cm, it is possible that disease risk saturates or is reduced above some species-specific size threshold (Fig. 2). Regardless of what the functional form of this relationship looks like across the entire range of colony sizes, the consistent importance of size across different host-disease pairs (effects ranging from 32 to 137% increase in disease risk) provides biological evidence for evaluating disease risk at the colony scale (also supported by simulation models [38]) rather than evaluating prevalence at the transect or site scale as is currently common practice. Such probabilistic models of colony health state could easily be integrated into a hierarchical framework to predict local or regional disease prevalence.

In addition to colony size, our findings suggest that shallow reefs with low herbivorous fish abundance, limited water motion, and located adjacent to watersheds with high fertilizer and pesticide runoff promote low levels of chronic disease year-round. We found that growth anomalies were more common in reefs with lower herbivorous fish abundance, potentially indicating indirect relationships with macroalgal cover that can harbor pathogens [39] or with more degraded reefs with high fishing intensity and associated impacts. Further, we found that growth anomalies were more likely to occur in shallow water, in reefs with low host density (likely because as colony sizes increase, host density decreases), and inside bays and other areas with low wave exposure (possibly because of longer residence times and associated concentration of pollutants). Growth anomalies in *P. compressa* and *P. lobata* were more common in locations with elevated levels of pesticide and fertilizer runoff from agriculture and golf courses (increasing disease risk by 23-80%, Table 4) and support previous research associating disease risk to nutrient enrichment, human populations, and poor water quality [17,40,41]. This research indicates that reducing coastal development and high-intensity land use activities such as farming and golf adjacent to semi-protected reefs would be an effective management approach to break transmission cycles of growth anomalies.

For tissue loss, we found that wave exposure, stream exposure, depth, and thermal stress were commonly associated with disease risk during interepidemic periods, but their precise effects sometimes differed across hosts. Tissue loss was consistently more common in shallow reefs with low thermal stress exposure (Table 5). The strong negative association with anomalously warm thermal stress in the summer for *P. compressa, P. lobata*, and *Porites* spp. models appears contrary to several previous studies on tissue loss in other regions [28,30,31,42]. Although there are no documented outbreaks of *Porites* tissue loss in Hawaii, this result may shed some light on why several tissue loss outbreaks in *Montipora* in Hawaii occurred in the winter [19, 33] compared with other regions where outbreaks occurred in the summer [30,31,43]. This result may suggest low temperature stress restricts the tissue loss pathogens’ growth rate, enabling the pathogen(s) to persist during interepidemic periods without causing an outbreak. Alternatively, disease risk could be highest in winter months because of a time lag between exposure to heat stress and increased host susceptibility to disease. We found that wave exposure and stream exposure were important risk factors for tissue loss as well, but the direction and magnitude of these relationships varied by host. For example, wave exposure was negatively associated with tissue loss in *M. capitata, Montipora* spp., and *P. evermanni* but positively associated with tissue loss in *P. lobata and Porites* spp. (Table 5). Similarly, stream exposure was negatively associated with *M. capitata* and *Montipora* spp. but positively associated with *P. compressa*. We did not find evidence that these opposing patterns were driven by differences in habitat associations across hosts (Figs. S2-3). One hypothesis that would explain this result is that water movement may be important for transmitting pathogens for some hosts (e.g., *P. lobata* and *Porites* spp.) but more stagnant water where pathogens can proliferate may be more likely to promote disease in other hosts (e.g., *M. capitata, Montipora* spp., and *P. evermanni*). Alternatively, the association with water movement may be related to water diffusion processes as colonies exposed to high water flow minimize the diffusion boundary layer [44] and could therefore limit exposure to pathogens in areas with high water movement and vice versa. Colony morphology can also affect water diffusion and pathogen exposure as branching morphologies (common in *Montipora* spp.) are more likely to retain water within their branches compared with massive colonies that tend to have smooth surfaces (common in *Porites* spp.). Similarly, stream exposure and potentially associated changes in salinity, sedimentation, and/or land-based pollutant runoff may promote disease occurrence in some hosts (e.g., *P. compressa*) and decrease disease risk in other hosts (e.g., *M. capitata* and *Montipora* spp.).

The variation in risk factors across host-disease pairs suggests that there may be different pathogens causing the same gross lesions in different hosts or that the same disease may arise in different hosts under different ecological conditions. Corals have a limited repertoire of responses to insult; therefore, disease lesions that are grossly identical, such as tissue loss, could be caused by different pathogens of the same general type (e.g., bacterial pathogens), which has been suggested in other studies [45]. Recent research supports this hypothesis as multiple bacterial pathogens have been identified for tissue loss in *Montipora capitata* in Hawaii [46]. Alternatively, the same causative agent(s) could be triggered to infect different species under different conditions. Our understanding of coral disease ecology is still limited but our results suggest that there may be specific ecological conditions that support or restrict the spread of specific diseases in each host. Thus, researchers should strongly consider how combining data for multiple species to assess disease risk may impact our understanding of factors that promote transmission. For example, combining species into a single model may be reasonable if the researchers acknowledge that the models will perform best for reefs that reflect the composition of species used in model development.

This study demonstrates the usefulness of the case-control design to study rare events in space and time using existing datasets with low probability of occurrence, but also provides a path forward for more efficiently collecting new data for future investigations. For example, for studies specifically focused on coral disease, we recommend complementing more traditional ecological assessments that characterize overall reef health, such as belt transect surveys and rugosity measurements, with survey efforts that maximize information on disease observations. Using experimental designs to collect data geared towards rare and/or clustered populations would allow for targetted investigations into the underlying mechanisms that drive disease processes. Therefore, for coral diseases, collecting more extensive information such as size, morphology, geographic location, clustering with nearby diseased colonies, severity of infection, area of affected tissue, etc. allows the use of modeling techniques to better explore the ecology of specific diseases. Such survey designs and associated analyses have already been developed and widely used in other fields such as epidemiology, plant pathology, and conservation biology (e.g., case-control design [47], cluster sampling [48, 49], line transect surveys [50]) and could be adapted for coral disease and other ecological studies. Rare occurrence or abundance data is very common in ecology and approaching ecological questions with this statistical design in mind will allow ecologists to more rigorously quantify ecological relationships for infrequent disturbances.

## Methods

### Coral health observations

We used coral health observations from the Hawaii Coral Disease Database (HICORDIS) [51] and additional surveys collected by the authors following the same methodologies used in HICORDIS (e.g., 10 x 1 m^2^ belt transects surveys) resulting in 362,366 coral health observations collected between 2004 and 2015. We separated the data by host (species or genus of interest) and disease type (growth anomalies or tissue loss) and used only observations with available data for all hypothesized risk factors (predictor variables). The final dataset excluded observations from the Northwestern Hawaiian Islands because several hypothesized risk factors were only available for the Main Hawaiian Islands. The number of disease and healthy colony observations we used for each host-disease pair are shown in Table 1.

### Ecological disease drivers

For each coral health observation, we collected biological, environmental, physical, and anthropogenic risk factors corresponding to the time and location of each observation (Tables 2-3). We selected risk factors based on relationships described in previous studies and based on personal observations (Table 2). After removing variables that were highly colinear (see *Testing for multicollinearity* below), our hypothesized risk factors were: (i) biological variables of colony size, host density, and herbivore fish abundance; (ii) environmental variables of anomalous temperature (Hot Snap), rainfall, chlorophyll-*a* (proxy for water quality/eutrophication), and irradiance; (iii) physical variables of depth, outside/inside of an embayment, acute and chronic wave exposure, and stream exposure (proxies for potential terrestrial runoff and retention); and (iv) anthropogenic variables of human population size and agriculture and golf course runoff. Variables reflect short-term (acute, pulse) or long-term (chronic, press) interactions or site characteristics (Table 3).

For each risk factor, we used survey data where recorded, or compiled externally sourced data from the vicinity (and within the watershed) of where the survey was conducted. We used measurements of colony size and depth that were recorded in the coral health surveys and calculated host density from the survey data. We extracted all other risk factors by survey location and by time (where applicable; i.e., acute interactions) using ArcGIS v 10.4. Measurements of rainfall were land-based, using gridded values from the land location nearest to the survey site that contained data within the appropriate watershed and month; human population size was similarly determined. For chlorophyll-*a*, wave exposure, irradiance, Hot Snaps, and herbivore fish abundance, we used data from the pixel containing the survey location when possible or else the nearest pixel with available data over the appropriate time period. However, we used herbivore fish abundance data compiled from only one year (2012-2013). We expect this data to adequately represent the spatial structure of herbivore fish abundance across the archipelago. Further, we expect year to year variation within the study period to be minimal, as the major drivers of herbivore fish dynamics – major disturbance events [52, 53] and protection status [54] – were minimized during the study period: there were no changes to marine protection status across the study region and only one major disturbance event (mass coral bleaching in 2015) at the end of the study period. We calculated agriculture and golf course runoff for each ocean pixel as the value from the nearest watershed decayed with distance from shore; therefore, we used the same technique as described above acquiring data from the pixel containing the survey location when possible or else the nearest pixel with available data. We manually determined whether a site was outside or inside of an embayment by visualizing the survey site locations relative to coastal boundaries downloaded from the State of Hawaii Office of Planning. We calculated stream exposure as the inverse planar distance between the site location and the nearest stream mouth within a given watershed. Higher stream exposure values indicate sites closer to stream mouths. For sites located in watersheds without streams, we set the stream exposure value to zero.

We used a different combination of risk factors for growth anomalies and tissue loss models because of hypothesized differences in their modes of transmission and etiologies. We included ten hypothesized risk factors in the growth anomalies models and eleven hypothesized risk factors in the tissue loss models; seven risk factors were common to both diseases (Table 2).

### Testing for multicollinearity

To test for multicollinearity among risk factors, we calculated the variance inflation factor. The variance inflation factor (VIF) is a ratio that quantifies the amount of multicollinearity among covariates in a regression model by dividing the variance of a model with multiple terms by the variance of a model with a single term. A VIF equal to one indicates a variable is not correlated with other variables, a value between one and five indicates moderate correlation, and a value above five indicates high correlation. We initially hypothesized that, in addition to the risk factors listed in Table 2, total coral density, wintertime thermal stress (Cold Snaps, Winter Condition), and coastal development should be included as risk factors for both growth anomalies and tissue loss models, and that an additional metric of urban runoff should be included in the growth anomalies model. However, these risk factors had VIF values greater than five indicating strong multicollinearity with other covariates (Fig. S4), so we excluded these risk factors from our candidate models.

### Logistic regression without the case-control design

We investigated the ecological drivers of each host-disease pair using logistic regression with all available coral health data (Tables 1-3) and assessed model predictive accuracies and true positive rates based on withheld portions of data (see below). The response variable in our logistic regression models described the colony observation health state as 0 for healthy and 1 for diseased. We included the hypothesized risk factors listed in Table 2 (combination dependent on disease type) for the full models with island and month as random effects and conducted backward model selection to find the best fit model based on the lowest Akaike Information Criterion (AIC). We included island and month as random effects to make the model results as comparable as possible with the case-control model results, as the case-control datasets were sub-sampled accounting for island and survey month. We conducted this analysis using the lme4 package [55] in R statistical software v 3.5.1. We used 80% of the data for model development and model selection, and the remaining 20% of data to quantify model predictive accuracy (proportion of total colony observations correctly identified as healthy and diseased) and true positive rate (proportion of disease colony observations correctly identified as diseased). We split the data using the sample function in base R, which selects data in proportion to the raw data (e.g., if the full data has a zero-to-one ratio of 90%-to-10%, then the sample will also contain 90% zeros and 10% ones).

To ensure that the randomly sampled training and testing data did not bias our results, we repeated our model development and selection and accuracy assessments 500 times. Therefore, for each host-disease pair, we split the full data into 500 paired training and testing datasets, conducted model selection within each training dataset, and then used the most frequently selected best fit model (Table S1) to predict health state on the 500 withheld test datasets. We compared predicted health state with observed health state to calculate accuracy and true positive rates within each withheld test dataset and report mean values across all 500 withheld test datasets.

### Logistic regression with the case-control design

For the logistic regression analysis with the case-control design, we followed the approach described above except instead of creating 500 randomly selected training and testing datasets from all available coral health data for each host-disease pair, we created 500 case-control datasets each with matching numbers of disease and healthy observations and randomly split those case-control datasets into paired training data and testing data (80% training/20% testing). To create each case-control dataset, we selected all disease colony observations for a given host-disease pair, and for each disease colony observation we randomly selected a healthy colony observation of the same host type (by species or genus) surveyed in the same month (but not necessarily year) and from the same island as the disease colony observation. We repeated this process to create 500 case-control datasets per host-disease pair that all had the same disease observations but varied in the paired healthy colony observations. We followed the same process of model development, selection, and testing as described above. Results for the frequency of model selection for the best fit model are available in Table S1.

## Acknowledgements

We thank the Fore-C research team for valuable feedback on analyses and the following funding sources for this research: NASA Roses Ecological Forecasting grant NNX17AI21G (MJD, SFH, JMC), NASA Earth and Space Science Fellowship (JMC, MJD), and Stanford University Woods Institute for the Environment Environmental Ventures Program (JMC). The scientific results and conclusions, as well as any views or opinions expressed herein, are those of the author and do not necessarily reflect the views of NOAA or the Department of Commerce.

## Author contributions

JMC, GA, and MJD designed experiments. JMC and SFH assembled data. JMC conducted the analyses and wrote the manuscript. JMC, GA, SFH, and MJD interpreted results and edited the manuscript.

---

**Figure S1:**
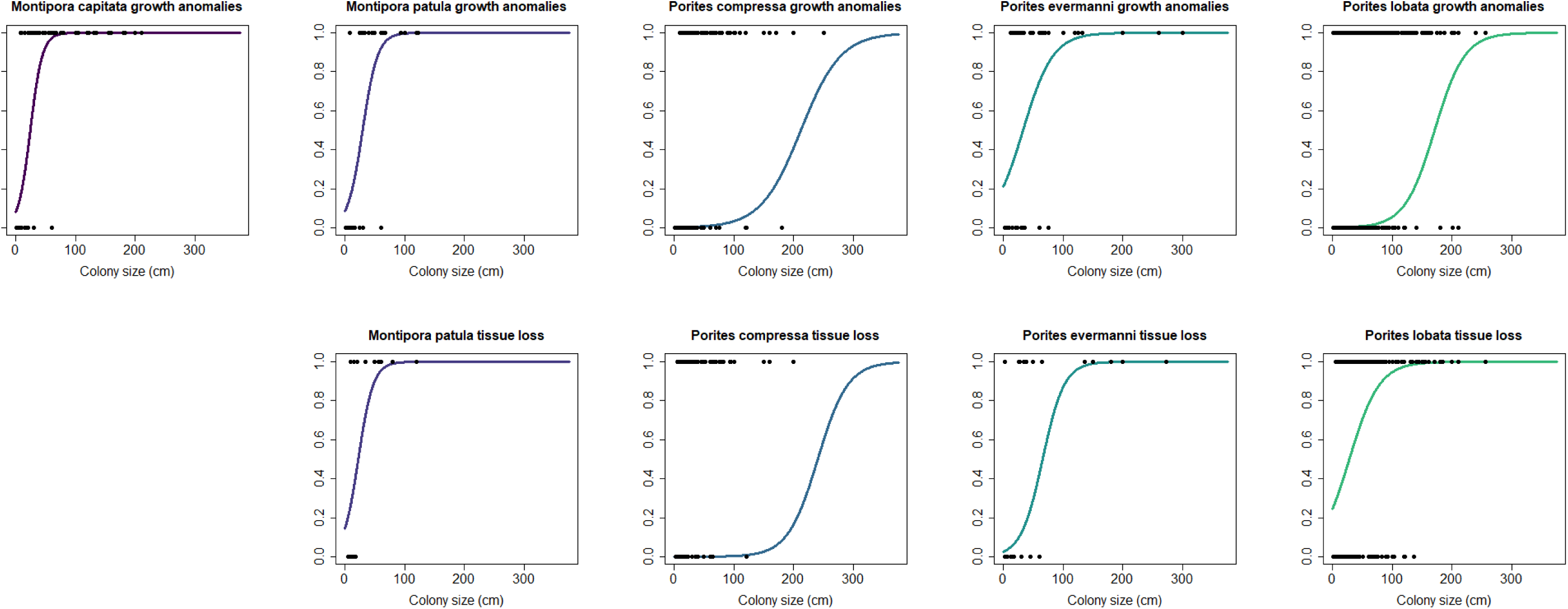
Disease risk as a function of colony size across host-disease pairs. Points indicate disease presence (P=1) and disease absence (P=0) from one random sample of a case control training dataset.

**Figure S2:**
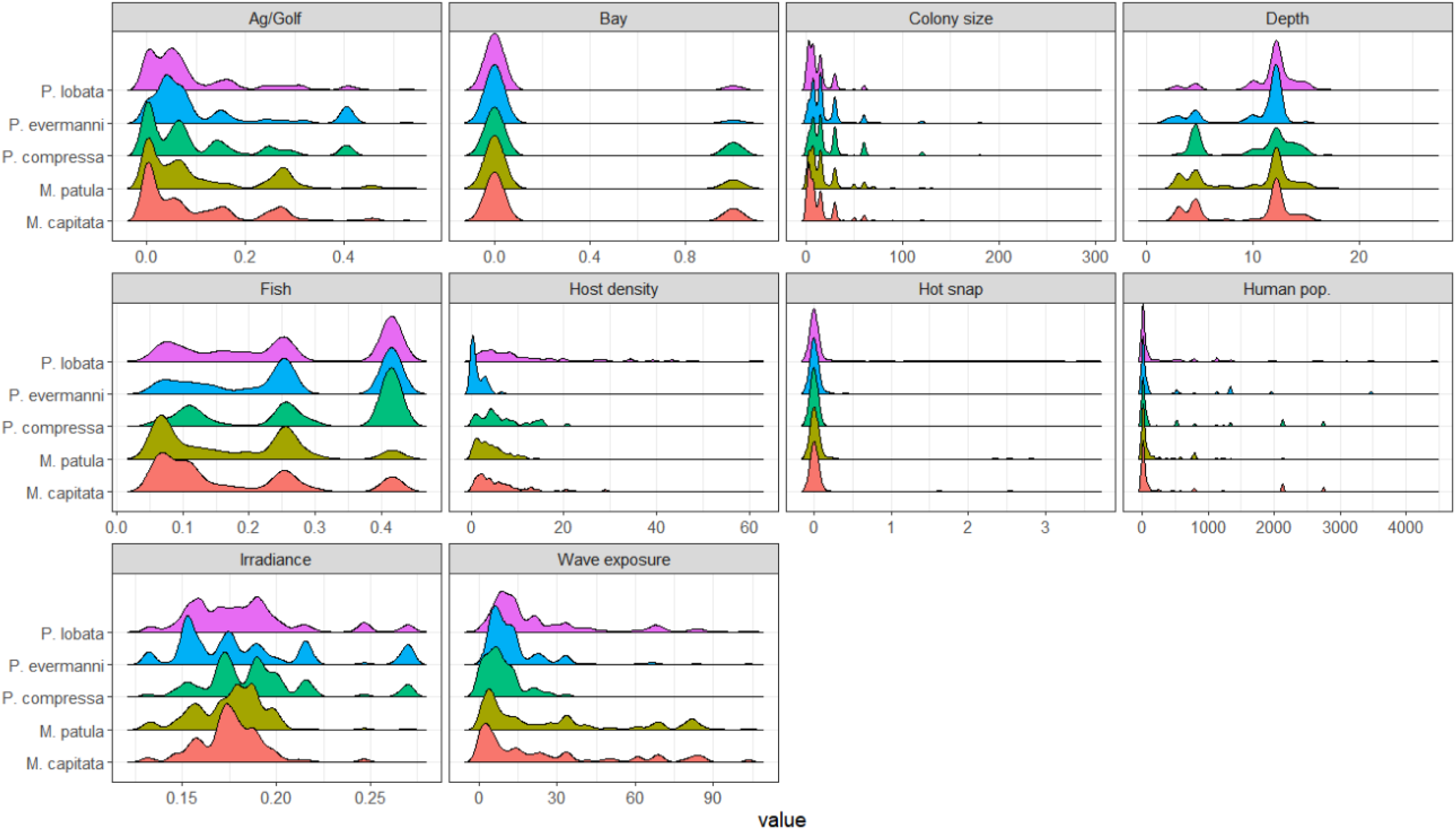
Distribution of risk factors (predictor variables) in growth anomaly models. Distribution of risk factors included in candidate growth anomaly models after removing observations with missing data. For the risk factor “Bay”, zero indicates outside a bay and one indicates inside a bay.

**Figure S3:**
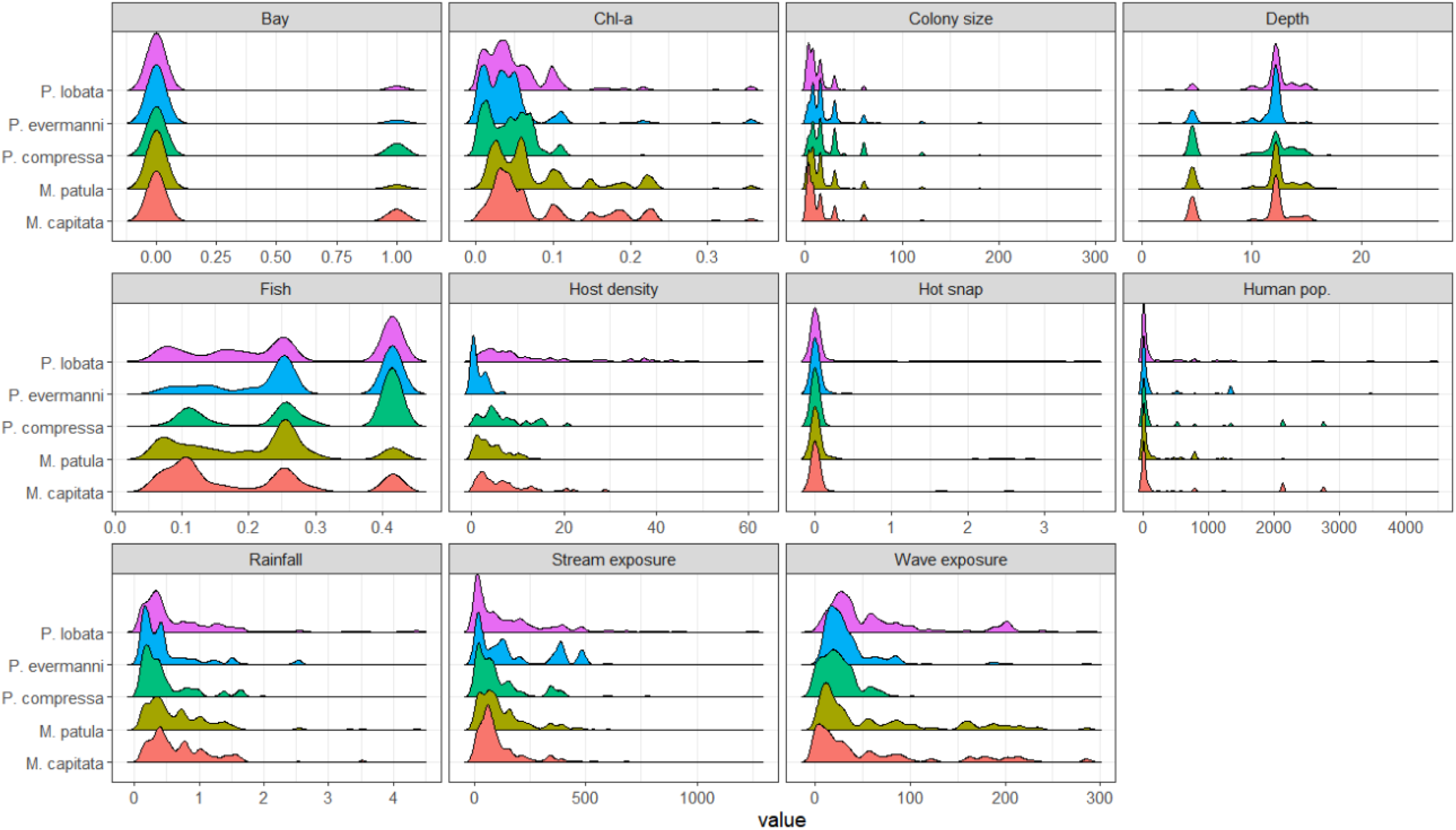
Distribution of risk factors (predictor variables) in tissue loss models. Distribution of risk factors included in candidate tissue loss models after removing observations with missing data. For the risk factor “Bay”, zero indicates outside a bay and one indicates inside a bay.

**Figure S4:**
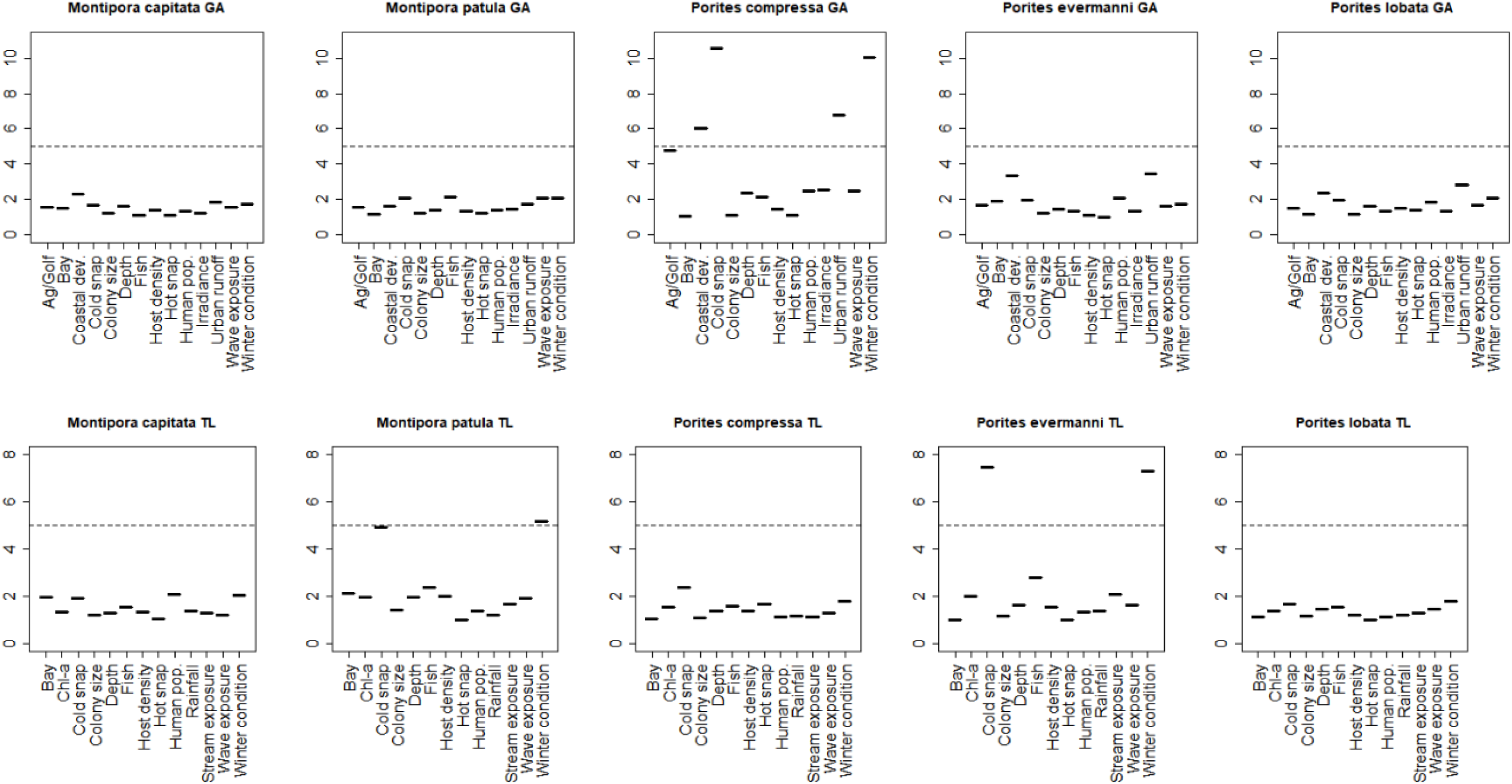
Variance inflation factor plots. Variance inflation factors for each predictor variable across host-disease pairs. Values greater than five (dashed horizontal line) indicates strong multicollinearity. For consistency, we removed predictor variables from all analyses if values exceeded five any host-disease pair.

**Table S1:** Number of times the best fit model was selected as the best model within the 500 training datasets. Disease refers to growth anomalies (GA) and tissue loss (TL).

| Disease | Species | Frequency of best fit model without case-control deign | Frequency of best fit model with case-control deign |
| --- | --- | --- | --- |
| GA | *Montipora capitata* | 497 | 76 |
| GA | *Montipora patula* | 125 | 73 |
| GA | *Montipora* spp | 176 | 69 |
| GA | *Porites compressa* | 212 | 113 |
| GA | *Porites evermanni* | 111 | 36 |
| GA | *Porites lobata* | 278 | 87 |
| GA | *Porites* spp | 354 | 83 |
| TL | *Montipora capitata* | 187 | 23 |
| TL | *Montipora patula* | 92 | 43 |
| TL | *Montipora* spp | 154 | 47 |
| TL | *Porites compressa* | 125 | 86 |
| TL | *Porites evermanni* | 216 | 30 |
| TL | *Porites lobata* | 474 | 90 |
| TL | *Porites* spp | 139 | 63 |

